# Mycobacterial infection of precision cut lung slices reveals that the type 1 interferon pathway is locally induced by Mycobacterium bovis but not M. tuberculosis in different cattle breeds

**DOI:** 10.1101/2021.04.16.440039

**Authors:** Aude Remot, Florence Carreras, Anthony Coupé, Émilie Doz-Deblauwe, ML Boschiroli, John A. Browne, Quentin Marquant, Delphyne Descamps, Fabienne Archer, Abrahma Aseffa, Pierre Germon, Stephen V. Gordon, Nathalie Winter

## Abstract

Tuberculosis exacts a terrible toll on human and animal health. While *Mycobacterium tuberculosis* (Mtb) is restricted to humans, *Mycobacterium bovis* (Mb) is present in a large range of mammalian hosts. In cattle, bovine TB (bTB) is a notifiable disease responsible for important economic losses in developed countries and underestimated zoonosis in the developing world. Early interactions that take place between mycobacteria and the lung tissue early after aerosol infection govern the outcome of the disease. In cattle, these early steps remain poorly characterized. The precision-cut lung slice (PCLS) model preserves the structure and cell diversity of the lung. We developed this model in cattle in order to study the early lung response to mycobacterial infection. *In situ* imaging of PCLS infected with fluorescent Mb revealed bacilli in the alveolar compartment, adjacent or inside alveolar macrophages (AMPs) and in close contact with pneumocytes. We analyzed the global transcriptional lung inflammation signature following infection of PCLS with Mb and Mtb in two French beef breeds: Blonde d’Aquitaine and Charolaise. Whereas lungs from the Blonde d’Aquitaine produced high levels of mediators of neutrophil and monocyte recruitment in response to infection, such signatures were not observed in the Charolaise in our study. In the Blonde d’Aquitaine lung, whereas the inflammatory response was highly induced by two Mb strains, AF2122 isolated from cattle in the UK and Mb3601 circulating in France, the response against two Mtb strains, H37Rv the reference laboratory strain and BTB1558 isolated from zebu in Ethiopia, was very low. Strikingly, the type I interferon pathway was only induced by Mb but not Mtb strains indicating that this pathway may be involved in mycobacterial virulence and host tropism. Hence, the PCLS model in cattle is a valuable tool to deepen our understanding of early interactions between lung host cells and mycobacteria. It revealed striking differences between cattle breeds and mycobacterial strains. This model could help deciphering biomarkers of resistance *versus* susceptibility to bTB in cattle as such information is still critically needed for bovine genetic selection programs and would greatly help the global effort to eradicate bTB.

## INTRODUCTION

Bovine tuberculosis (bTB) caused by *Mycobacterium bovis* (Mb) remains one of the most challenging infections to control in cattle. Because of its zoonotic nature, this pathogen and associated notifiable disease in cattle are under strict surveillance and regulation in the European Union. When bTB cases are detected through surveillance, culling of these reactor cattle is mandatory. In spite of intensive eradication campaigns, bTB is still prevalent in European cattle [1; 2] and has significant economical, social and environmental implications. Since 2001, France is an officially bTB free country a status that was achieved through costly surveillance programs. However, each year, around one hundred Mb foci of infection are identified [3] with certain geographical areas showing a constant rise in disease prevalence since 2004.

bTB eradication is an unmet priority that faces two major difficulties: the persistence of undetected infected animals in herds because of lack of diagnostic senstivity, and the risk of transmission from infected sources [4]. Moreover, the poor understanding of bTB pathophysiology in cattle and the lack of correlates of protection are substantial knowledge gaps that must be resolved so as to better tackle the disease (DISCONTOOLS, https://www.discontools.eu/).

Both Mb and *M. tuberculosis* (Mtb) belong to the same genetic complex. Mtb is responsible for tuberculosis (TB) in humans which displays similar features with bTB. It is estimated that one third of the global human population are latently infected with Mtb which kills 1.4 million people each year [5]. Despite the high degree of identity that Mtb and Mb share both at the genetic level as well as during the infection process, the two pathogens display distinct tropism and virulence depending on the host. While Mb is highly virulent and pathogenic for cattle and a range of other mammals, Mtb is restricted to sustain in humans. Experimental infection of cattle with the widely used Mtb laboratory strain H37Rv, which was genome sequenced in 1998 [6], shows strong attenuation as compared to Mb [7; 8]. However, natural infection of cattle with Mtb has been reported, and the strain Mtb BTB1558 was once such case, isolated from a zebu bull in Ethiopia [9; 10]. In comparison to the original UK Mb strain AF2122/97, the first genome sequenced Mb isolate [11; 12], the Mtb strain BTB1558 diplayed much lower virulence in European cattle [13].

The Mb strains that circulate in France today are phylogenetically distant from the UK Mb reference strain. While AF2122 belongs to the European 1 clonal complex [14], the European 3 clonal complex is widespread in France, [15]. The Eu3 genetic cluster is composed of field strains that share the SB0120 spoligotype with the attenuated Bacillus Calmette Guérin vaccine strain [16; 17]. In our study, we used Mb3601 as the reprensentative strain of this widespread French cluster. Originally, Mb3601 was isolated from a tracheobronchial lymph node of an infected bovine in a bTB highly enzoonotic area in France [16]. However, despite widespread circulation in its origine area, nothing is known today of the pathophysiology of Mb3601 infection.

Indeed, greater knowledge is available on Mtb infection process and disease development both in humans and mouse models as compared to Mb infection in cattle. With both mycobacteria, the alveolar macrophage (AMP) is the frontline cell that first presents the first niche for mycobacteria entering the lung, and the role of the AMP in early-stage infection is well established [8]. Both Mtb and Mb have established their lifestyle in AMPs: they can escape its bactericidal mechanisms and multiply within this niche. During the infection process, bacilli disseminate to different anatomical sites and establish new infection foci both in the lungs and secondary lymphoid organs [18; 19]. During Mtb infection, lung epithelial cells also play key roles in the host defense (reviewed in [20; 21; 22]). Type II pneumocytes are infected by Mtb [23] and produce pro-inflammatory cytokines which augment the AMP innate resistance mechanisms [24]. The role of Type II pneumocytes during Mb infection in cattle is not well known. Also, most of the available knowledge on the role of bovine macrophages (MPs) during Mb infection comes from studies conducted with monocytes sampled from blood and derived as MPs during *in vitro* culture [25].

In our study, we wanted to investigate the bovine innate response following Mb or Mtb infection in a preserved lung environment to allow the resident lung cells to interact with bacilli and crosstalk. Precision cut lung slices (PCLS) are an experimental model in which resident lung cell types are preserved and remain alive for at least one week [26]. The tissue architecture and the interactions between the different cells are maintained. PCLS have already been validated for the study of various respiratory pathogens [26; 27; 28]. In chicken PCLS, mononuclear cells are highly motile and actively phagocytic [29]. This model is well designed to study complex interactions taking place early after the host-pathogen encounter. During Mb infection in cattle, important differences in production of key proinflammatory cytokines such as IFNγ or TNFα by peripheral blood mononuclear cells are observed depending on the clinical status of the animal. Interestingly, such differences are observed at early time points [30] indicating that the innate phase of the host reponse is key to the establishment of the pathological outcome of the infection.

Therefore the PCLS model is ideally suited to investigate early host-pathogen interactions in the bovine lung during Mb infection, and may help to find clues to the impact of the innate response on the outcome of infection. This model, that fully mimicks the early environment of the bacillus entering the lung (as compared to monocyte-derived MPs) may also aid in understanding the molecular basis of mycobacterial host preference [31]. To this end, we decided to compare four mycobacterial strains: two Mtb species, namely the Mtb H37Rv reference strain for human TB and the cattle-derived Mtb BTB1558, and two Mb species namely Mb AF2122 as representative of the EU1 clonal complex and Mb3601 as the hallmark EU3 strain. Since the host genetic background also has profound impact on the outcome of bTB disease [32], we decided to compare PCLS from two prevalent beef breeds in France, Charolaise and Blonde d’Aquitaine, and conducted a thorough characterization of the lung responses to Mb and Mtb during *ex vivo* infection. PCLS allowed us to decipher important differences in the transcriptomic and cytokine profile during the innate response to infection, depending both on the breed, i.e. between Blonde d’Aquitaine and Charolaise cows, and on the mycobacterial species, i.e. between Mtb and Mb.

## MATERIALS AND METHODS

### Animal tissue sampling

Lungs from fifteen Blonde d’Aquitaine and nine Charolaise cows were collected post-mortem at a commercial abattoir. Animals were between three- and eleven-years-old and originated from eight different French departments where no recent bTB outbreak had been notified (Figure S1). No ethical committee approval was necessary as no animal underwent any experimental procedure. After slaughter by professionals following the regulatory guidelines from the abattoir, the lungs from each cow were systematically inspected by veterinary services at the abattoir. The origin of each animal was controlled and its sanitary status was recorded on its individual passport: animals were certified free of bTB, leucosis, brucellosis, and infectious bovine rhinotracheitis.

### Bacterial strains and growth conditions

Strains Mb AF2122/97 and Mb MB3601 had previosuly been isolated from infected cows in Great Britain and France, respectively [12; 15]. The Mb3601-EGFP fluorescent strain were derived by electroporation with an integrative plasmid expressing EGFP and selected with Hygromycin B (50 μg/mL) (Sigma, USA) as described in previously [33]. Mtb BTB1558 had been previosuly isolated from a zebu bull in Ethiopia [13]. Bacteria were grown in Middlebrook 7H9 broth (Difco, UK) supplemented with 10% BBLTM Middlebrook ADC (Albumin- Dextrose- Catalase, BD, USA) and 0.05% Tween 80 (Sigma-Aldrich, St Louis, USA). At mid-log phase, bacteria were harvested, aliquoted, and stored at −80°C. Batches titers were determined by plating serial dilutions on Middlebrook 7H11 agar supplemented with 10% OADC (Oleic acid- Albumin-Dextrose-Catalase, BD, USA), with 0.5% glycerol or 4.16 g/L sodium pyruvate (Sigma, USA) added for Mtb or Mb strains, respectively. Plates were incubated at 37°C for 3-4 weeks (H37Rv, BTB558 and AF2122) and up to 6 weeks for Mb3601 before CFUs numeration. Inocula were prepared from one frozen aliquot (titer determined by CFU numeration) that was thawed in 7H9 medium without glycerol and incubated overnight at 37°C. After centrifugation 10 min at 3000 x g, concentration was adjusted to 10^6^ cfu/mL in RPMI medium.

### Obtention and infection of Precision-Cut Lung Slices (PCLS)

PCLS were obtained from fresh lungs using a tissue slicer MD 6000 (Alabama Research and Development). For each animal, the right accessory lobe was filled via the bronchus with RPMI containing 1.5% low melting point (LMP) agarose (Invitrogen) warmed at 39°C. After 20 min at 4°C, solidified lung tissue was cut in 1.5 cm slices with a scalpel. A 0.8 mm diameter punch was used to obtain biopsies that were placed in the microtome device of the Krumdieck apparatus, filled with cold PBS, and 100 μM thick PCLS were cut. One PCLS was introduced in each well of a P24 well plate (Nunc), one mL of RPMI 1640 (Gibco) supplemented with 10% heat inactivated Fetal Calf Serum (FCS, Gibco), 2 mM L-Glutamine (Gibco) and PANTA™ Antibiotic Mixture (polymyxin B, amphotericin B, nalidixic acid, trimethoprim and azlocillin; Becton Dickinson) was added to the well and the plate was incubated at 37°C with 5% CO_2_. Medium was changed every 30 min during the first 2 h to remove all traces of LMP agarose. Twenty four hours later, after the last medium change, the ciliary activity was observed under a microscope to ensure tissue viability.

PCLS were infected two days with 10^5^ CFU of Mb or Mtb strains. As indicated, PCLS were either fixed in formalin for imaging or lyzed with a Precellys in lysing matrix D tubes in 800 μL Tri-reagent for RNA extraction. The bacillary load of each strain present in the PCLS was compared after transfer of PCLS to a new plate 1 day after infection (dpi), two washes in 1 mL of PBS, and homogenization in 1 mL of PBS in lysing matrix D tubes (MP Biomedicals) with a Precellys (Ozyme). To determine CFUs, serial dilutions were plated as described above.

### Alveolar macrophages (AMPs)

To harvest AMPs from Blonde d’Aquitaine cows, broncho-alveolar lavages (BAL) were performed on the left basilar lobe of the lung at a local abattoir after culling of the animal. The lobe was filled with 2 x 500mL of cold PBS containing 2 mM EDTA (Sigma-Aldrich). After massage, the BAL was collected and transported at 4°C to the laboratory. BAL was filtered with a 100 μm cell strainer (Falcon) and centrifuged for 10 min at 300 x g. Cells were washed in RPMI medium supplemented with 10% heat inactivated fetal calf serum (Gibco), 2 mM L-Glutamine (Gibco) and PANTA™ Antibiotic Mixture. 10^7^ BAL cells per mL were suspended in 90% FCS and 10% DMSO (Sigma-Aldrich) and cryopreserved in liquid nitrogen. One day before infection, BAL cells were thawed at 37°C, washed in complete RPMI medium, and transferred to a 75 cm² culture flask with a ventilated cap. After 2 h at 37°C 5% CO_2_, non adherent cells were removed, and adherent AMPs were incubated 2 x 10min at 4°C with 10 mL of cold PBS to detach and enumerate them in a Malassez chamber. 5 x 10^5^ AMPs /well were distributed in a P24 well-plate and incubated overnight at 37°C 5% CO_2_. Medium was changed once and AMPs were infected with Mb3601 or Mtb H37Rv at a MOI of 1. At 6 h and 24 h pi, supernatants were filtered through a 0.2 μm filter and cells were lyzed in 800 μL of Tri-reagent for RNA extraction. MOI was checked by CFU determination 24 h after infection.

### Cell supernatant collection and lactate deshydrogenase (LDH) assay

In order to evaluate cytotoxicity, supernatants from infected PCLS or AMPs were passed through a 0.2 μm filter at indicated time points and cells were lysed in 1 mL of lysis buffer (5mM EDTA, 150 mM NaCl, 50 mM Tris-HCl, Triton 1%, pH 7,4), containing anti-proteases (Roche), in lysing matrix D tube, with a Precellys apparatus. Homogenates were clarified by centrifugation 10 min at 10000 x g, filtered through 0.2 μm and collected on microplates. Cytotoxicity of infection in PCLS was assessed using the “Non-Radioactive Cytotoxicity Assay” kit (Promega) according to the manufacturer’s instructions. The cytotoxicity was calculated as: cytotoxicity (%) = (OD_490_ of LDH in the supernatant)/(OD_490_ of LDH in the supernatant + OD_490_ of LDH in the PCLS homogenates) x 100.

### Immunohistochemistry on PCLS

Infected PCLS were fixed 24 h at 4°C with 4% of formalin, then transfered to a 48-wells culture plate in PBS. All steps described below were done under gentle agitation at room temperature (RT). PCLS were incubated 2 h with 100 μL of PBS, 0.25% Triton X-100, 10% horse serum for permeabilization and saturation (saturation buffer). They were incubated overnight at 4°C with primary Ab (anti-bovine MHCII clone MCA5655 from BioRad, anti-bovine pancytokeratine clone BM4068 from Acris) diluted in saturation buffer. PCLS were washed 4 times with 300 μL of PBS (2x 5 min, then 2x 10 min), then incubacted 3 h with fluorescent-conjugated secondary antibodies diluted in saturation buffer (Goat anti-mouse IgG1-APC and Goat anti-mouse IgG2a A555 from Invitrogen). PCLS were washed 4 times with 300 μL of PBS (2x 5 min, then 2 x 10 min), and transfered on coverslides which were mounted with Fluoromount-G™ Mounting Medium, containing DAPI (Invitrogen) and sealed with transparent nail polish. Z-stack imaging was performed at x 63 enlargment with a confocal microscope (LEICA) and analyzed with LAS software.

### Quantification of cytokines and chemokines released by PCLS and AMPs

Cytokine and chemokine levels produced by PCLS after 2 dpi were assessed in a Multiplex assay in supernatants (dilution 1:2) with MILLIPLEX® Bovine Cytokine/Chemokine Panel 1 (BCYT1-33K-PX15, Merck) according to the manufacturer’s instructions. IFNγ, IL-1α, IL-1β, IL-4, IL-6, IL-8 (CXCL8), IL-10, IL-17A, IL-36RA (IL-1F5), IP-10 (CXCL10), MCP-1 (CCL2), MIP-1α (CCL3), MIP-1β (CCL4), TNFα, & VEGF-A were measured. Data were acquired using a MagPix instrument (Luminex) and analyzed with Bio-Plex Manager software (Bio-Rad). IL-8 was out of range in the Multiplex, so we performed a sandwich ELISA with the following references: Goat anti Bovine Interleukin-8 Ab AHP2817, Recombinant Bovine Interleukin-8 PBP039 and Goat anti Bovine Interleukin-8 Ab conjugated to biotin AHP2817B (all from Bio-Rad, protocol according to the manufacturer’s intructions).

### RNA extraction and gene expression analysis

Total RNA from two pooled PCLS were extracted using a MagMAX™-96 Total RNA isolation kit (ThermoFisher). For AMPs we used the Nucleospin RNA isolation kit (Macherey Nagel). After DNase treatment (ThermoFisher or Macherey Nagel) mRNAs were reverse transcribed with iScript™ Reverse Transcriptase mix (Biorad) according to the manufacturer’s instructions. Primers (Eurogenetec; Supplementary Table S1) were validated using a serially diluted pool of cDNA mix obtained from bovine lung, lymph nodes, blood and bone marrow, with a LightCycler® 480 Real-Time PCR System (Roche). Gene expression was then assessed with the BioMark HD (Fluidigm) in 96 x 96 well IFC plate, according to the manufacturer’s instructions. The annealing temperature was 60°C. Data were analyzed with Fluidigm RealTime PCR software to determine the cycle threshold (Ct) values. Messenger RNA (mRNA) expression was normalized to the mean expression of three housekeeping genes (*PPIA*, *GAPDH*, *ACTB*) to obtain the ΔCt value. For each animal, values from infected PCLS were normalized to the uninfected PCLS gene expression (ΔΔCt value, and Relative Quantity = 2 ^− ΔΔCt^). Principal Component Analysis (PCA) were performed using ΔΔCt values in R studio (Version 1.1.456 © 2009-2018 RStudio, PBC), using the FactoMineR packages (version R 3.5.3).

### Statistical analysis

Individual data, and the median and interquartile range are presented in the figures, exept for Figure 2 where the mean and standard deviation are presented. Statistical analyses were performed with Prism 6.0 software (GraphPad). Analyzes were performed on data from two to six independent experiments, with 2-way ANOVA or Wilcoxon non-parametric tests for paired samples used. Represented p-values were: \**p* < 0.05; \*\**p* < 0.01; \*\*\**p* < 0.001.

## SUPPLEMENTARY INFORMATION

The Supplementary Material for this article can be found online.

## RESULTS

### *Ex vivo* infection with mycobacteria of live bovine lung tissue in PCLS allows bacilli uptake by AMPs and their recruitment to alveoli

Early events of bTB pathophysiology in the bovine lung remain poorly defined due to the complexity of biocontained experimental infection in large animals. Since PCLS have been used to study viral respiratory infections in the bovine [26], we decided to use this model to assess early events taking place following entry of Mb into the lung. We infected bovine PCLS obtained *ex vivo* with the four mycobacterial strains: Mb AF2122, Mb3601, Mtb H37Rv or BTB1558.

We first monitored tissue cytotoxicity at 1 and 2 days post infection (dpi) using a lactate deshydrogenase (LDH) release assay. The mean percentage of cytotoxicity remained below 10% and no difference was observed between infected and non-infected PCLS (Fig. 1A). The ciliary activity from the PCLS bronchial cells monitored every day under a light microscope remained vigourous and stable after infection (data not shown). We calibrated our model and inocula to use 10^5^ CFUs for each of the four different strains. We analyzed CFUs still present in PCLS 24 h later and observed an equivalent 1 log decrease for all strains (Fig. 1B). This indicated equivalent infection by all strains, allowing them to be directly compared. Therefore, with similar bacterial load and excellent tissue viability in all experimental conditions, we validated PCLS as a model to study early events taking place in the bovine lung after infection with mycobacteria.

**Figure 1:**
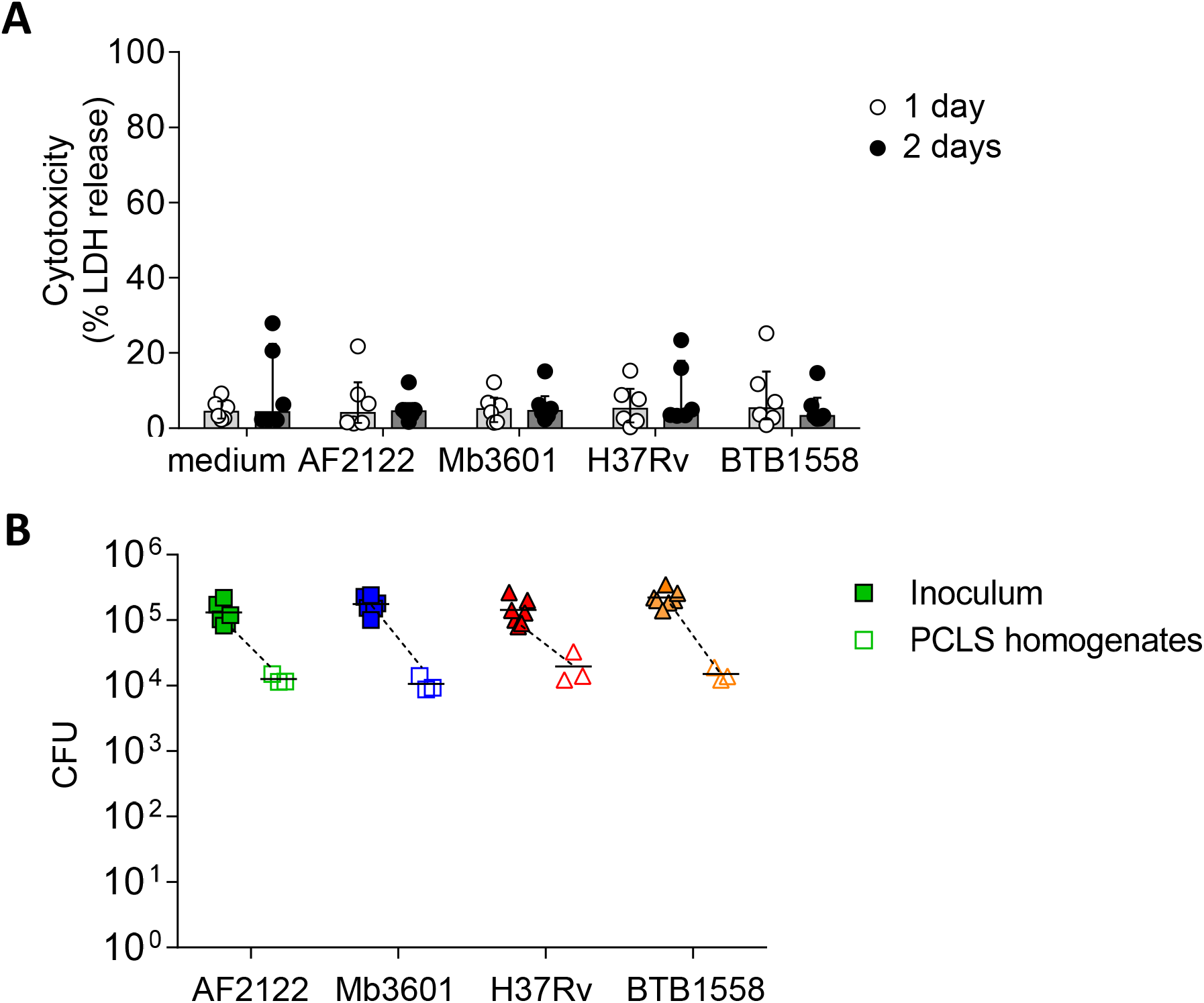
PCLS infection with four different Mb or Mtb strains does not induce lung tissue cytotoxicity and equivalent numbers of bacilli are recovered 24 h post-infection. **(A)** PCLS prepared from Blonde d’Aquitaine lungs post-mortem were infected with 10^5^ cfu of two Mb strains (AF2122 or Mb3601) or two Mtb strains (H37Rv or BTB1558). After 1 and 2 days post infection, PCLS supernatants were harvested and tissue was homogenized. Lactate deshydrogenase (LDH) was measured in both compartments using the “Non-Radioactive Cytotoxicity Assay” kit. Cytotoxicity was determined as (%) = (O.D.490nm LDH in supernatant) / (O.D.490nm LDH in supernatant + O.D.490nm LDH in PCLS homogenates) x 100. Individual data and the median and interquartile range in each group are presented (n=6 animals, from 6 independent experiments). **(B)** 24h post infection, PCLS were washed and homogenized to recover bacilli. Inoculum and PCLS homogenates were serially diluted and plated with CFUs numerated after 3-6 weeks incubation. Individual data and the mean in each group are presented (n=6 independent inocula prepared, PCLS homogenates data represent the mean of technical duplicates from n=3 animals, from 3 independent experiments).

In order to visualize interactions taking place between bacilli and lung cells, we infected PCLS with a fluorescent version of the Mb3601 strain, and at 1 and 2 dpi we analyzed the cells by *in situ* immunohistochemistry. The lung structure was visualized by DAPI and pancytokeratine staining and we used confocal microscopy to image 10-15μm sections and localize Mb3601-EGFP (Fig. 2A). We observed Mb in 27 ± 3 % of PCLS alveoli (Fig. 2A and 2B) and almost always in close contact with large MHC-II positive AMPs. Bacilli were localized outside AMPs in 76 ± 2 % observations and resided intracellularly in AMPs in 24 ± 2 % (Fig. 2A and 2C and Supplementary video). Interestingly, the number of AMPs per alveoli differed upon bacilli presence or absence (Fig. 2D). In uninfected PCLS, lung alveoli generally contained one AMP (data not shown). However, in Mb infected PCLS, we either observed no AMPs in 66 ± 2 % of alveoli or one AMP in 33 ± 2 % of alveoli in the absence of any Mb. On the contrary, the number of AMPs significantly increased in alveoli where at least one Mb was observed (Fig. 2D, *p*<0.001). The number of AMPs varied among infected alveoli with 24 ± 9 % containing one AMP, 52 ± 6% containing 2 or 3 AMPs and 9 ± 4 % containing more than 4 AMPs. Such observations indicated that during the 2 days of infection, AMPs were recruited from one alveoli to the other in response to signals linked to Mb infection. In conclusion, even though Mb infection was performed *ex vivo*, bacilli were observed in the alveoli, close or inside their target host cell i.e. the AMP. Moreover, the PCLS model was physiological enough to allow AMPs to crawl in response to signals linked to bacilli entry.

**Figure 2:**
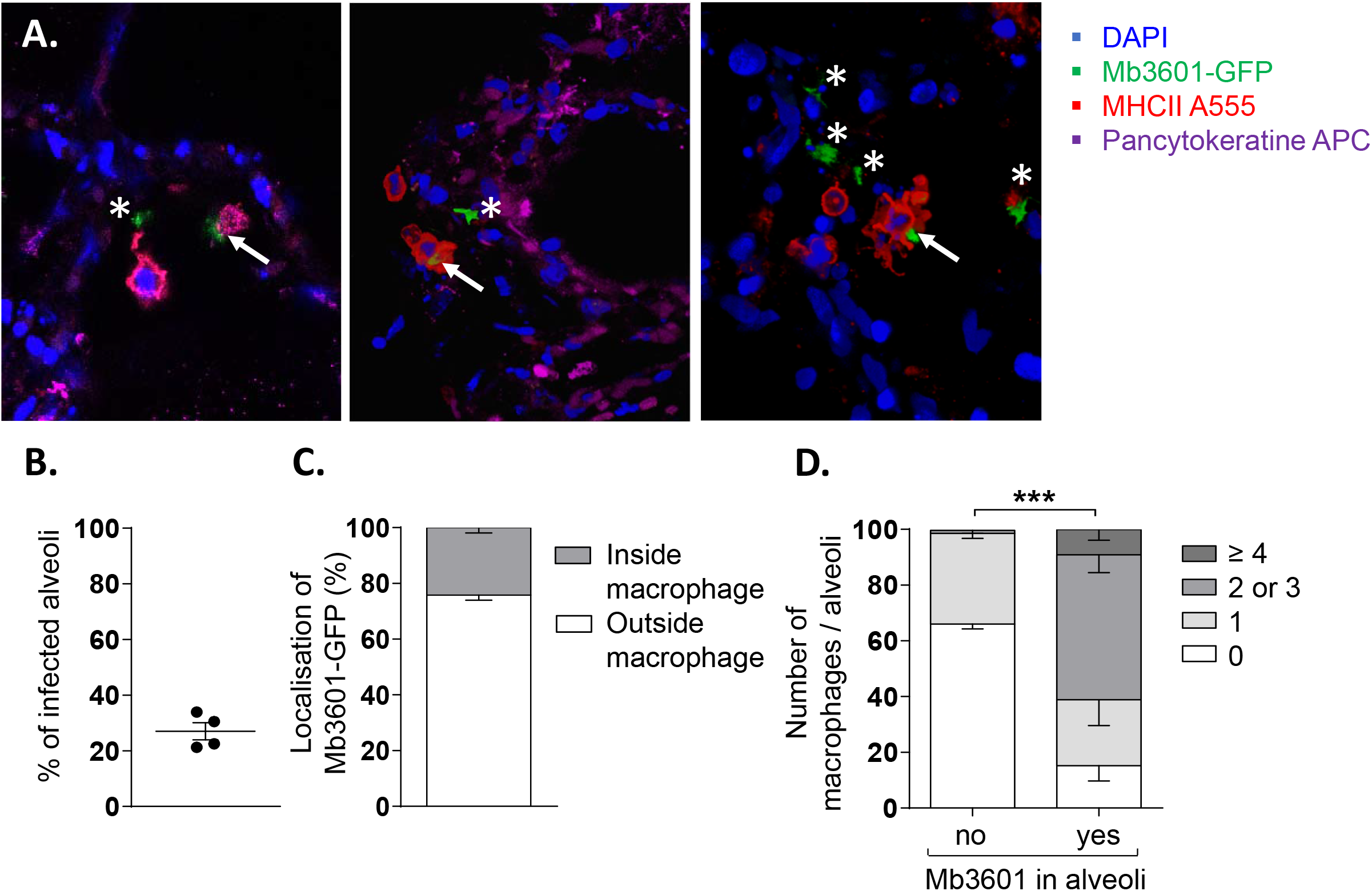
Mb3601 is internalized by AMPs in the preserved lung structure from PCLS and infected alveoli contain higher numbers of AMPs as compared to non-infected alveoli. PCLS were infected with 10^5^ CFUs of the green fluorescent Mb3601-GFP recombinant strain and fixed 2 days later. After labelling with anti-pancytokeratine (APC, magenta) and anti-MHCII antibodies (Alexa 555, red), PCLS were mounted with Fluoromount-G™ Mounting Medium containing DAPI (blue) and analyzed under a Leica confocal microscope **(A)**; 3D images were analyzed with Leica LAS software. Z-stack imaging was performed at x63 enlargment (10-15μm of thickness, step size of 0.5-1μm). White asterisks indicate extracellular bacilli and white arrows indicate bacili inside MHC-II^pos^ AMPs. **(B)** Graph represents the percentage of infected alveoli per PCLS among the 55 to 80 alveoli that were observed under the microscope (n=4 PCLS from two different Blonde d’Aquitaine cattle) **(C)** Stack histogram of the mean percentage +/− SEM of intra or extracellular bacilli among a minimum of 15 infected alveoli that were observed (N=4 PCLS) **(D)** The number of MHC-II^pos^ AMPs per alveoli was counted in infected or non-infected alveoli. The data presented as % are the mean +/− SEM of n=4 PCLS from two different Blonde d’Aquitaine cattle. Between 55 and 80 alveoli were observed to obtain these data (two way ANOVA, *** *p*<0.001)

### The lung response to mycobacterial infection vastly differs between Blonde d’Aquitaine and Charolaise cows

Two bovine beef breeds are widely used in France: Blonde d’Aquitaine and Charolaise. We decided to compare how these two breeds respond to mycobacterial infection, using our PCLS system. We measured 15 cytokines and chemokines secreted by the lung tissue at 2 dpi with the four mycobacterial strains and performed a principal component analysis (PCA). As depicted in Fig. 3A, PCA revealed important differences in the immune response of the lung tissue between the two breeds. Group samples clearly plotted appart and their ellipses showed either a small overlay (AF2122 and Mb360A) or no overlay at all (H37Rv). Results for the BTB1558 group showed less clustering of samples due to higher individual variations. We then extracted total RNA from PCLS after 1 or 2 dpi and analyzed the expression of 96 genes related to innate immunity and inflammation (see full list in supplementary Table 1). RT-qPCR data were normalized and expressed as fold change compared to uninfected PCLS control for each cow. Gene expression was higher 2 days after infection as compared to 1 (data not shown). We therefore decided to focus our analysis on this 2 dpi time point. Remarkably, the transcriptomic signature induced by infection was very low for the Charolaise breed, whichever mycobacterial strain was used, which explains the clustering of Charolaise samples (Fig. 3B). Increasing the inocumlum in the Charolaise PCLS up to 5 x 10^6^ CFU did not induce gene expression (Fig. S2). The response of the lung tissue to mycobacterial infection in Blonde d’Aquitaine was very different as compared to Charolaise, as revealed by a PCA (Fig. 3B). Whereas in PCLS from Charolaise gene expression from infected and non-infected controls clustered, in PCLS from Blonde d’Aquitaine, gene expression levels were significantly more dispersed after infection as compared to controls (Fig 3B). We compared individual gene expression between the two breeds for a number of genes. For instance, both the *CXCL2* chemokine, and the mycobacteria receptor syndexin 4 *SDC4*, were significantly upregulated after PCLS infection with AF2122, Mb3601 or H37Rv in Blonde d’Aquitaine, but not in Charolaise (Fig. 3C). Altogether, our data revealed important differences in the early lung response to mycobacterial infection, depending on the breed of the animals, that could be measured both at the gene expression and protein production level in the PCLS system.

**Figure 3:**
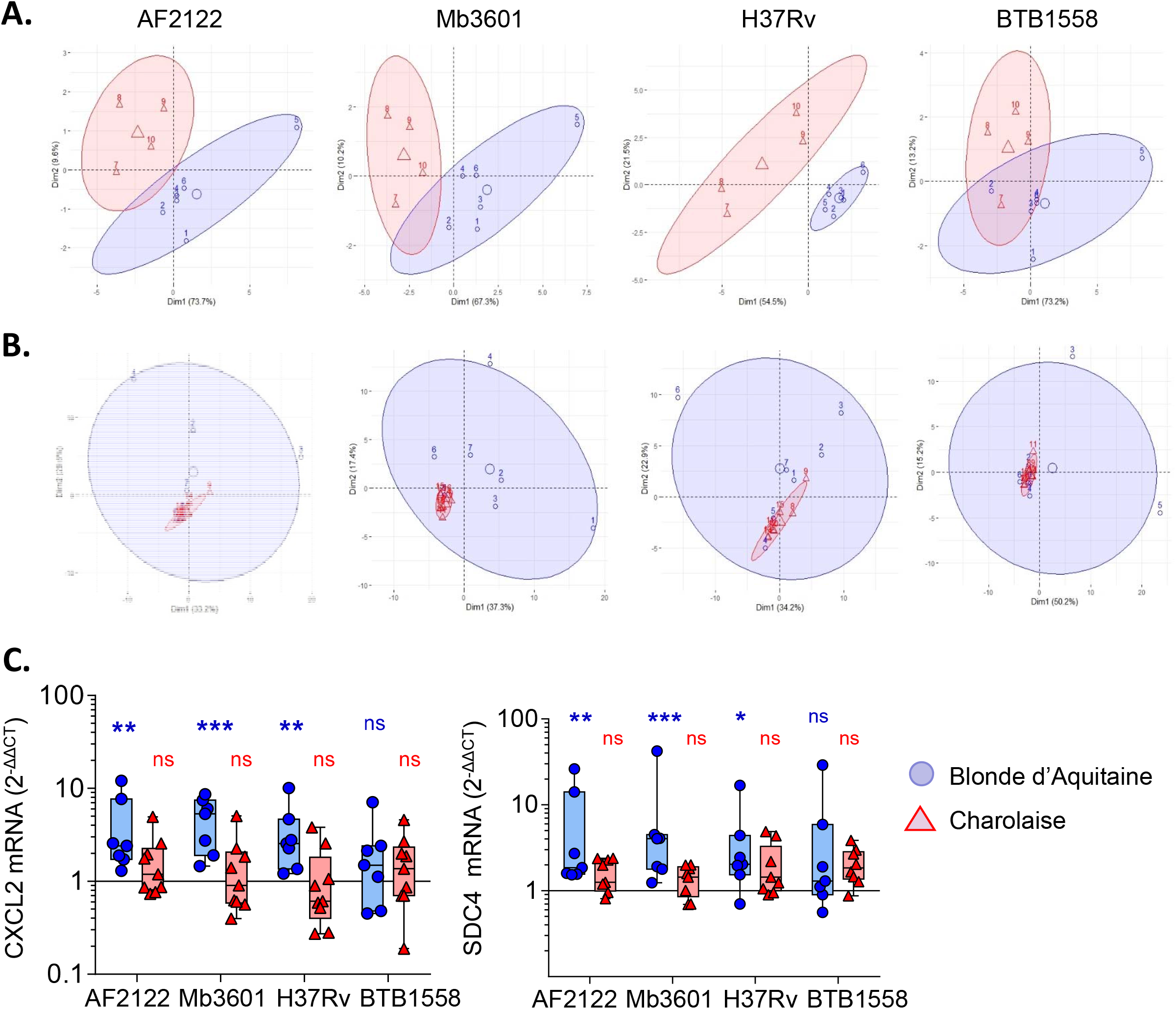
Principal Component Analysis (PCA) of inflammatory lung tissue signature reveals differences between two beef cattle breeds after 2 days of infection by Mb or Mtb. **(A)** Fifteen cytokines and chemokines were measured in PCLS supernatants from Blonde d’Aquitaine or Charolaise cows 2 days after infection with four different mycobacterial strains. Raw data were used to run PCA in R studio. Individual data are shown (n=4 for Charolaise, red; n=6 for Blonde d’Aquitaine, blue). Ellipses represent a confidence range of 90%. **(B)** PCA were built from expression data of 96 genes (2^−ΔΔCt^) obtained from PCLS total RNA extracted 2 days after infection. Individual data are shown (n=9 for Charolaise, red; n=7 for Blonde d’Aquitaine, blue). Ellipses represent a confidence range of 90%. **(C)** Two examples of differentially expressed genes. Individual data and the median and interquartile range in each group are presented (n=7 Blonde d’Aquitaine and n=9 Charolaise) * *p*<0.05. ** *p*<0.01. *** *p*<0.001. Two way ANOVA test.

### The overall inflammation signature in the lung tissue is triggered more efficiently by *M. bovis* than *M. tuberculosis*

We then focused our analysis on Blonde d’Aquitaine to determine how the lung tissue responded to different mycobacterial strains. We analyzed 15 cytokines and chemokines produced in the PCLS supernatants 2 days following infection. No IL-4 was detected and production of TNFα, IL-36RA, IL-10, VEGFA or MCP-1 was not different between infected PCLS and controls (Figure S3A). We observed that *ex vivo* infection of PCLS with mycobacteria triggered an inflammatory response that contrasted between the strains (Fig. 4A). At the protein level, the Mtb strain BTB1558 induced the most heterogenous response and, due to high individual variation, differences in chemokine/cytokine production between infected PCLS and controls only reached statistical signficance for MIP-1a (CCL3) and IL-8 (Fig. 4A). These two inflammatory mediators were also strongly induced by all strains. IL-17A, IL-1β and IFNγ were efficiently induced by mycobacterial infection and no significant difference was observed between Mtb and Mb. By contrast, IL-6 and IL-1α were significantly induced after Mb but not Mtb infection and IL-8 production was also significantly higher after Mb than Mtb infection (Fig 4A). The only strain able to induce significant production of MIP-1b was Mb3601. We then analyzed the inflammatory transcriptomic signature using a panel of 17 genes involved in monocyte/macrophage and neutrophil recruitment (Fig. 4B). A number of these genes was significantly upregulated upon PCLS infection even though significant differences were not always reached due to inter-individual variation. Remarkably, Mb3601 induced the strongest inflammatory response, with 5 out of 17 genes significantly upregulated as compared to non-infected controls. Focusing on chemokines involved in neutrophil recruitment, we observed that *CXCL2* expression was induced by all strains -except BTB1558- whereas *CXCL1*, *CXCL5* and *CXCL8* were only upregulated by Mb3601 (Fig. 4B and 4C). *IL-6* expression was also high after Mb3601 infection. Therefore, *ex vivo* infection of PCLS efficiently triggered signals involved in monocyte and neutrophil recruitment. Infection by Mb strains, and more specifically the Mb3601 strain circulating in France, triggered inflammation in the bovine lung more efficiently than Mtb.

**Figure 4:**
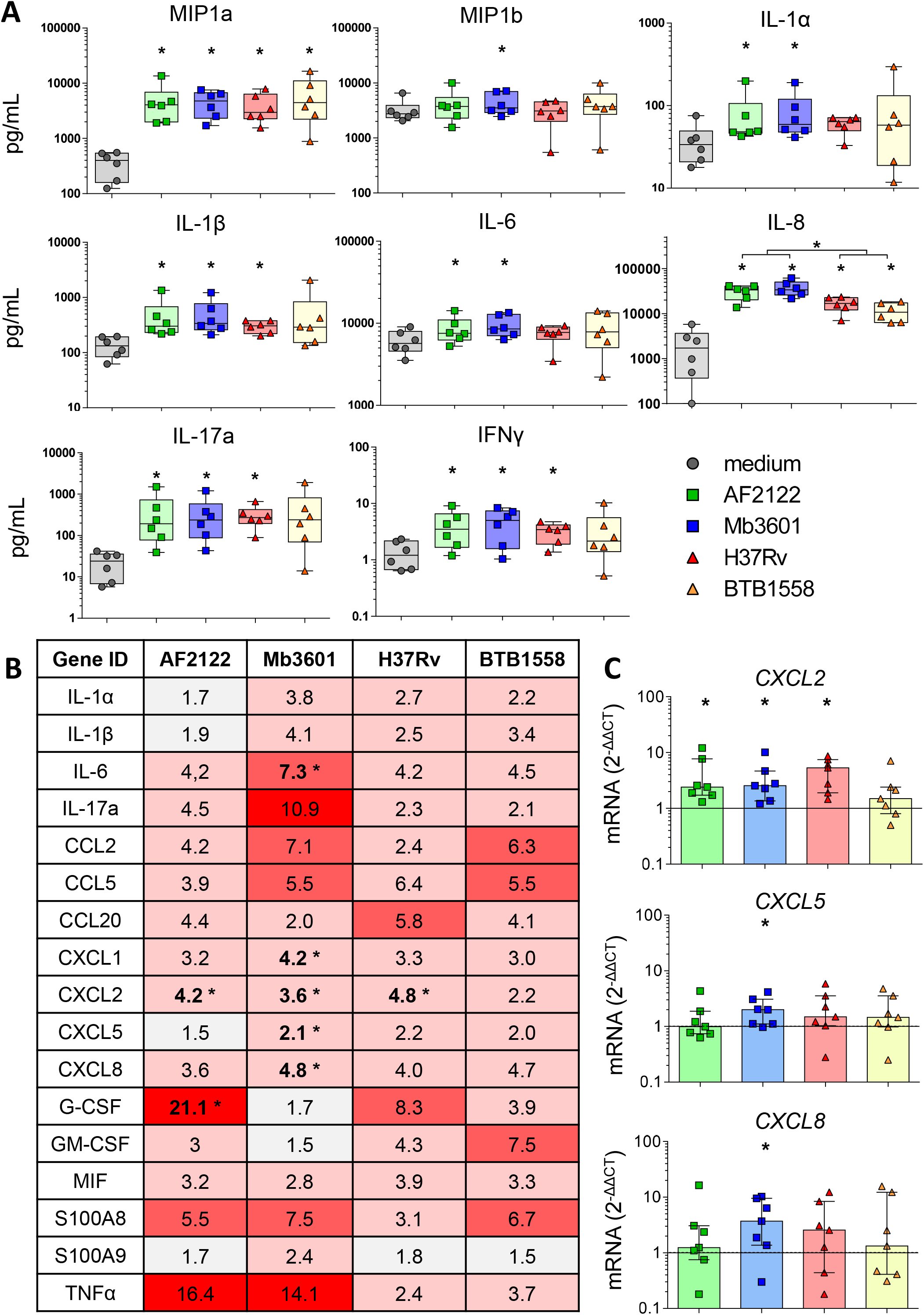
The lung inflammatory neutrophil and monocyte recruitment signature induced by infection in PCLS from Blonde d’Aquitaine cows is more efficiently triggered by M. bovis than M. tuberculosis. **(A)** Cytokine and chemokine levels were measured in PCLS supernatant by Multiplex ELISA two days after infection with two Mb or two Mtb strains. Individual data and the median and interquartile range in each group are presented (n=6 cows). **(B)** Table of the mean of fold change (2^−ddCT^) for each group (n=7 cows) of 17 major genes involved in neutrophil and monocyte recruitment and inflammation. The graduated red box coloring represents levels of gene expression, and asterisks mark significant differences compared to non-infected controls. **(C)***CXCL2*, *CXCL5* and *CXCL8* gene expression at 2 days post infection. Individual data and the median and interquartile range in each group are presented (n=7 cows). (B-C) * *p*<0.05 (Wilcoxon non parametric test).

### The type I interferon pathway is induced in the bovine lung by infection with *M. bovis* but not *M. tuberculosis*

Because in humans and mouse models, susceptibility to mycobacterial infection and disease progression is driven by type I IFN [34; 35; 36], we decided to compare induction of this pathway by Mtb and Mb strains in bovine lung tissue. We measured expression of different genes involved in the type I IFN pathway in Blonde d’Aquitaine PCLS infected by the four mycobacterial strains (Fig. 5). Gene expression of both *IFNβ* and the *IFNAR1* receptor were significantly increased after Mb but not Mtb infection (Fig. 5A and 5C). Similarly, the major IFN stimulated genes (ISG) *MX1*, *OAS1*, *ISG15* and *CXCL10*, were induced only after Mb infection (Fig. 5A and 5C) and this difference was also detected at the protein level for CXCL10 (Fig. 5A and 5B). Therefore we observed induction of a number of genes of the type I IFN pathway, recapitulated in Fig 5D, after infection with Mb but not Mtb strains. Strikingly, strain Mb3601 was the highest inducer of this pathway in the lung from Blonde d’Aquitaine cows.

**Figure 5:**
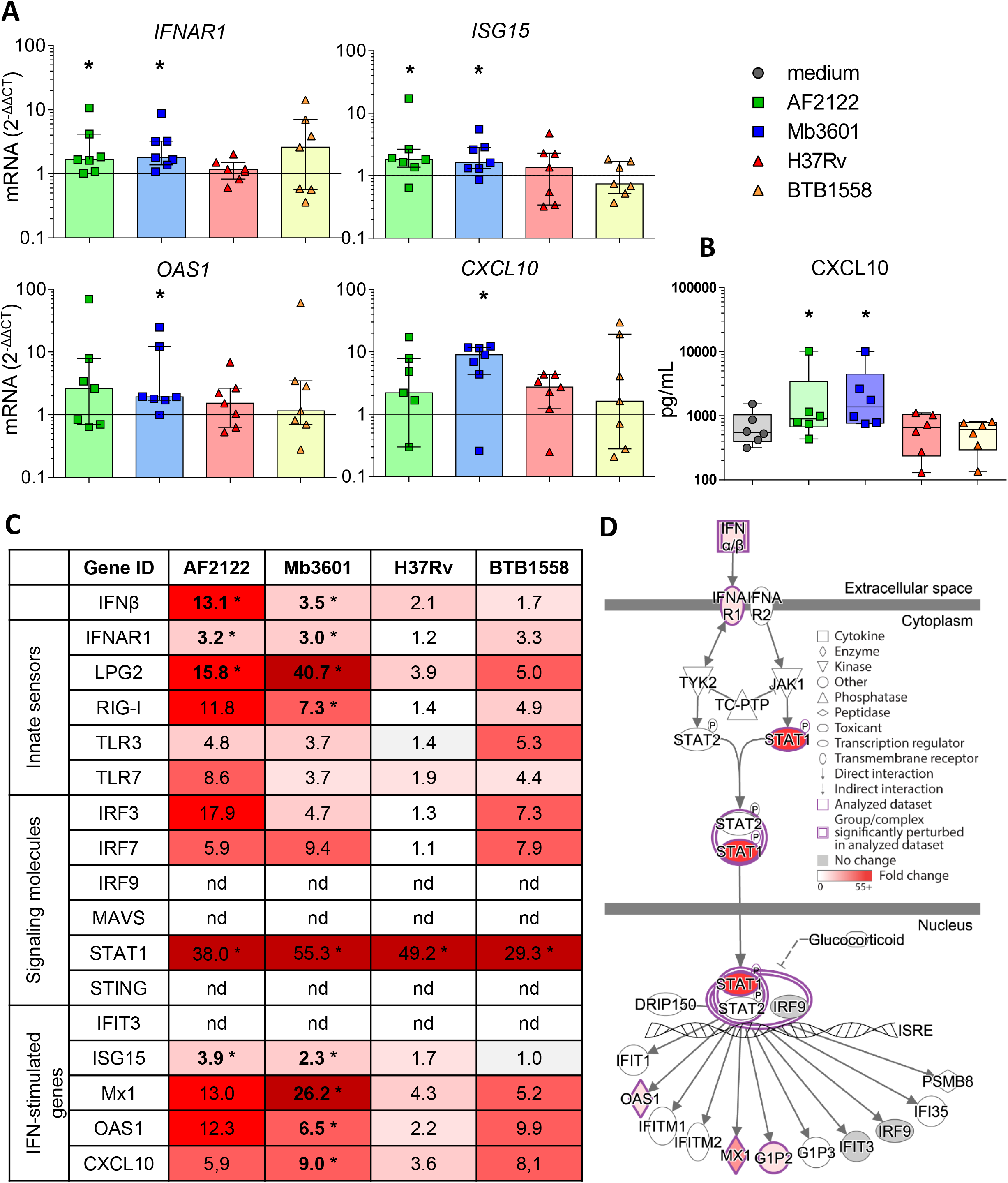
Mb but not Mtb infection in the lung tissue from Blonde d’Aquitaine cows induces the type I interferon pathway. PCLS were infected as described in Figure 1. **(A)** *IFNAR*, ISG15*, CXCL10* and *OAS1* gene expression at 2 dpi. Individual data and the median and interquartile range in each group are presented (n=7) **(B)** CXCL10 protein level was measured in PCLS supernatant at 2 dpi. Individual data and the median and interquartile range in each group are presented (n=6). **(C)** The table represents the mean of fold change (2^−ddCT^) for each group (n=7) of major genes involved in type I interferon pathway. Graduated red box coloring are for higher gene expression and asterisks mark significant differences compared to uninfected PCLS. nd=not detected. **(D)** Ingenuity Pathway Analysis drawing of the Type I interferon pathway under IFNAR in the Mb3601 group. Graduated red box coloring are for higher gene expression. (A, B and C) * *p*<0.05 (Wilcoxon non parametric test).

Because AMPs are the most promiment host cell interacting with Mb [8], which we also observed in PCLS (Fig. 2), we next decided to decipher if AMPs contributed to induction of the type I IFN pathway after Mb3601 or H37Rv infection. One day after infection of AMPs with these two strains similar bacterial levels were recovered (data not shown). At 6 hours post infection no cell cytotoxicity was observed and we analyzed expression of genes from the type 1 IFN pathway at this early time point. While we did not observe differences in *IFNAR1*, *IRF3*, *STAT1* nor *ISG15* expression induced by the two strains (Fig. 6A), *IFNβ*, *LPG2*, *RIG1* and *OAS1* were significantly induced after infection with Mb3601 but not H37Rv (Fig. 6C and 6D). Regarding *MX1*, the same trend was observed although statistical significance was not reached (Fig. 6D, *p*=0.07).

**Figure 6:**
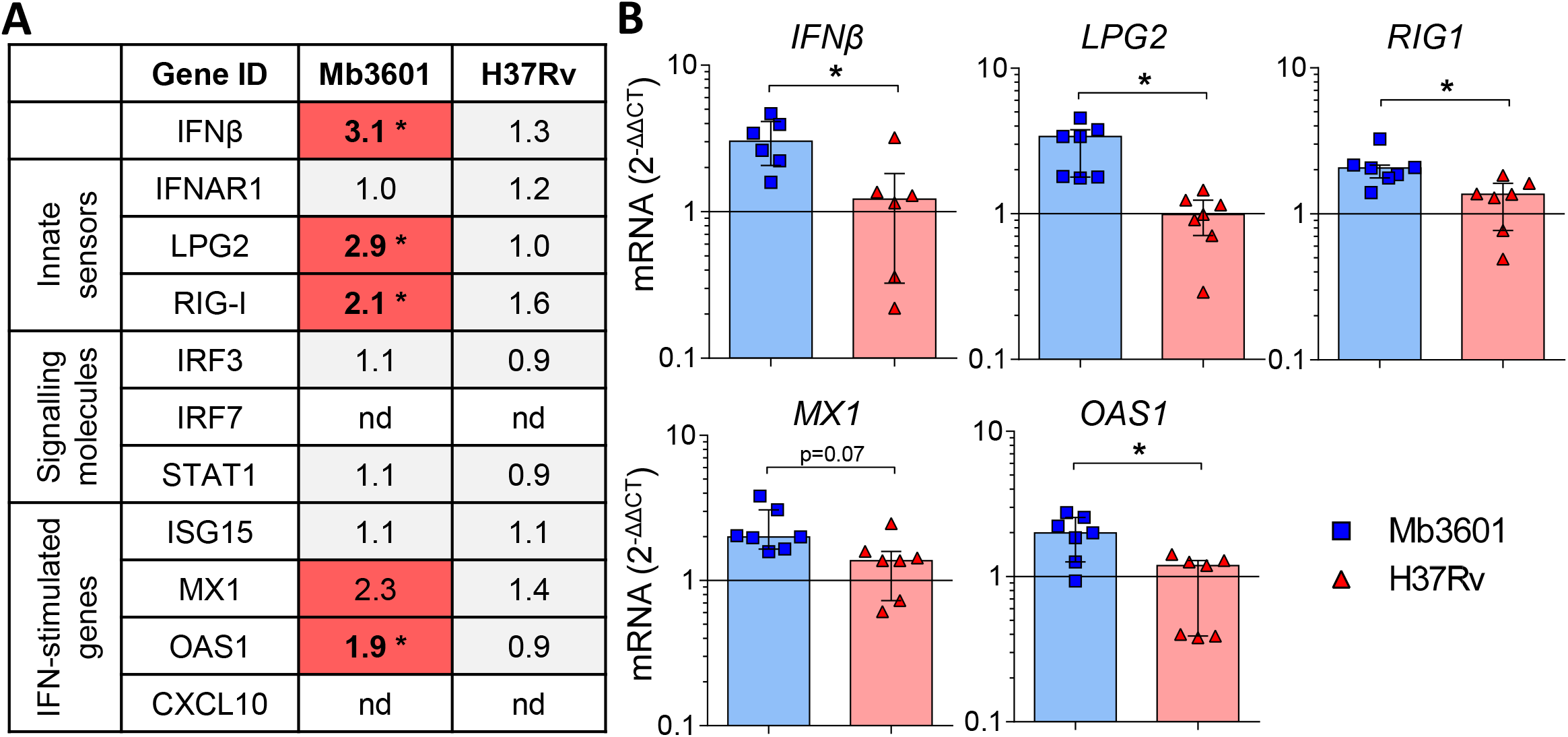
Alveolar macrophages from Blonde d’Aquitaine contribute to the type I IFN signature in lung induced by Mb infection. AMPs from Blonde d’Aquitaine lungs were infected with 10^5^ cfu of Mb3601 or Mtb H37Rv. 6 h later mRNA was extracted from and expression of major genes from the type 1 IFN pathway was analyzed **(A)** Mean fold change (2^−ddCT^) of gene expression normalized to three house keeping genes was calculated in each group (n=7). Graduated red box coloring represents gene expression and asterisks mark significant differences compared to non infected controls (nd=not detected). **(B)** *IFNβ*, *LPG2*, *RIG1*, *MX1* and *OAS1* gene expression in AMPs was analysed by RT-qPCR at 6h post infection. Individual data and the median and interquartile range in each group are presented (n=7). **(A, B)** * *p*<0.05 (Wilcoxon non parametric test).

Interestingly, while CXCL10 was detected both at the mRNA and protein level in PCLS infected with Mb (Fig. 5), we did not detect expression of this gene by AMPs in our analysis. Altogether these results demonstrate that AMPs globally contribute to the type I IFN pathway in the lung after Mb infection, although other cells present in PCLS may also specifically induce some genes, such as *CXCL10* or *IRF7* for example (Fig. 5C and 6A).

## DISCUSSION

The lung is the main organ targeted by Mb infection in cattle [37] and early interactions between the different lung cell types and the bacillus that govern the pathophysiology of the disease need to be better understood. In this study we used PCLS for the first time to monitor the early bovine lung response to Mb infection, and validated this model as a means to measure the local innate respoense at the protein and mRNA level. A main advantage of PCLS is conservation of the complex lung tissue both in structure and diversity of cell types. After infection with mycobacteria, ciliary activity of bronchial cells was maintained. AMP main function is to patroll the lung, crawling in and between alveoli, they sensed, chemotaxed, and phagocytosed debris or inhaled bacterial [38]. We observed increased numbers of AMPs in alveoli where Mb was present, indicating AMP mobility inside the tissue. In chicken, PCLS allowed observation of MPs movement and phagocytosis [29]. The AMP is well established as the main host cell for Mtb infection in humans [39] and Mb infection in cattle [40]. Accordingly, in PCLS we observed Mb inside AMPs in 20% of infected alveoli. We sometimes observed several bacilli inside one AMP. Although Mb is able to replicate inside this hostile cell, it is difficult to know if this observation was due to bacillary multiplication or phagocytosis of several bacilli. This issue would need live imaging of PCLS to follow the fate of fluorescent Mb, an approach which remains challenging under BSL3 conditions.

In uninfected PCLS, we observed generally one AMP for 2-3 alveoli (Fig S4, in good correlation with Neupane et al., [38]). After Mb infection, we observed several AMPs inside the same alveolus in 50% cases. Moreover, when the alveoli contained more than four AMPs, they were in closed contact. Multinucleated giant cells are formed by fusion of several MPs and are a hallmark of TB pathophysiology. It has recently been demonstrated that after infection of human or bovine blood derived MPs by Mb or Mtb, only Mb was able to induce the formation of multinucleated cells [41]. Although 2 dpi we did not observe formation of such cells in PCLS, it would be interesting to analyze if such events could be detected after longer infection periods.

One other advantage of our model is the preserved diversity of lung cell composition. PCLS contain type I and type II pneumocytes, endothelial cells, and bronchial cells (Fig S4) and also produce key molecules like surfactant which has an established role in Mtb uptake [42]. Mtb is also capable of invading type II alveolar epithelial cells [23] that play important roles in host defense [20; 21; 22]. In our study, we did not observe intraepithelial Mb, but specific labelling of bovine epithelial cells would be required to investigate interactions between bovine lung pneumocytes and Mb in more detail. However, as we observed that infected AMPs were in close contact with epithelial cells in PCLS, this model will allow a more refined analysis of the crosstalk between AMPs and pneumocytes during Mb infection [24].

One limitation of the PCLS model is the lack of recruitment of immune cells from circulating blood. During mycobacterial infection, in response to local signals, a variety of immune cells are recruited to the infection site to form the mature granuloma that constrains bacillary multiplication. How this reponse is orchestrated at the level of the lung tissue in cattle remains poorly established. Neutrophils, together with other innate cells such as macrophages, γδ-T lymphocytes and natural killer cells, were recently identified as key immune cells in early containment of infection [43] and development of early lesions [44]. Moreover, humans regularly exposed to Mtb, or cattle exposed to Mb, do not always develop signs of infection, i.e. remain negative in IFNg-release assay or skin testing. In humans, such resistance to infection through successful elimation of bacilli could be mediated by neutrophils [45]. Similarly, in cattle experimentally infected with Mb, some contact animals resist infection while others develop lesions due to productive infection [46]. It is possible that neutrophils could also play an important role in early elimination of Mb in cattle [43]. Immune signals involved in early recruitment of neutrophils to the lung after entry of Mb need to be better understood in cattle. It is known that epithelial cells secrete, among other cytokines and chemokines, MIP1 and CXCL8 that attract MPs and neutrophils to the site of infection. Interetingly we measured important differences in production of such mediators by PCLS in response to different strains of mycobacteria that could be linked to variable virulence. Although one cattle Type II pneumocyte cell line has been described [47], such transformed cells are less physiologically relevant than primary cells. Thus, PCLS could help understanding the early orchestration of the local inflammatory response in the lung in response to mycobacterial infection.

Resistance to bTB is linked to the host genetics. Zebu breeds (*Bos indicus*) have been reported to be more resistant than Bos taurus derived breeds to bTB disease [48]. Our results with PCLS, as a physiological model of the early lung response to infection, demonstrated striking differences between Blonde d’Aquitaine and Charolaise emphasizing the importance of the host genetics in response to Mb. It is not known whether the stronger inflammatory response of the Blonde d’Aquitaine tissue is associated with greater sensitivity or resistance to Mb infection. While robust immunological responses are associated with increased pathology at the level of the animal [30], at the cellular level, blood derived MPs from animals with greater resistance to bTB (and that kill BCG more efficiently than cells from susceptible animals) produce higher levels of the pro-inflammatory mediators iNOS, IL-1β, TNFα, MIP1 and MIP3 [49]. Although genetic selection of cattle would greatly complement bTB management and surveillance programs to control and ultimately eradicate the disease, especially in countries with the highest burden [50; 51], biomarkers to evaluate resistance or susceptibility of cattle to Mb infection are critically missing. Some genomic regions and candidate genes have been identified in Holstein Friesian cows, the most common dairy breed [52] and not surprisingly, these candidates are often involved in inflammation. A genomic region on chromosome 23, containing genes involved in the TNFα/NFκ-B signalling pathway, was strongly associated with host susceptibility to bTB infection [53]. However, large within-breed analyses of Charolaise, Limousine, and Holstein-Friesian cattle identified 38 SNPs and 64 QTL regions associated with bTB susceptibility to infection [54]. The genotyping of 1966 Holstein-Friesian dairy cows that were positive by skin test and either did, or did not, habour visible bTB lesions, together with their skin-test negative matched controls led to the conclusion that these variable phenotypes following Mb exposure were governed by distinct and overlapping genetic variants [55]. Thus, variation in the pathology of Mb seems to be controlled by a large number of loci and a combination of small effects. Similar conclusions were drawn from genetic studies of human tuberculosis [56]. In areas where Mb is highly prevalent, recurrent exposure to Mb may also imprint the bovine genome, and epigenetics could also contribute to the immune response in certain breeds. In France, the Nouvelle Aquitaine region accounted for 80% of Mb outbreaks last year. Interestingly, Blonde d’Aquitaine breed is very abundant in this area (Fig. 7). Together with another very abundant beef breed in this region, Limousine, they contribute to most bTB outbreaks in Nouvelle Aquitaire (bovine tuberculosis national reference laboratory communication). Future comparisons with Limousine would be interesting. In our study, Blonde d’Aquitaine or Charolaise cows were sampled from eight different French departments, none with recurrent Mb outbreaks, rendering previous exposure to Mb unlikely. Moreover breeding management was similar for the two breeds, as far as we could ascertain, suggesting that exposure to environment and possible wildlife sources would be comparable. We nevertheless observed striking differences in the early lung response to Mb infection between these two breeds, pointing to possible control of Mb infection at the genetic or epigenetic level. Whether one cattle breed is more susceptible to bTB than others remains an open question that deserves future studies, with more consequent animal sampling to better sustain our observations. We furthermore believe that the PCLS model could greatly contribute to unravelling the role of tissue-level protective responses that would in turn reveal important biomarkers.

**Figure 7:**
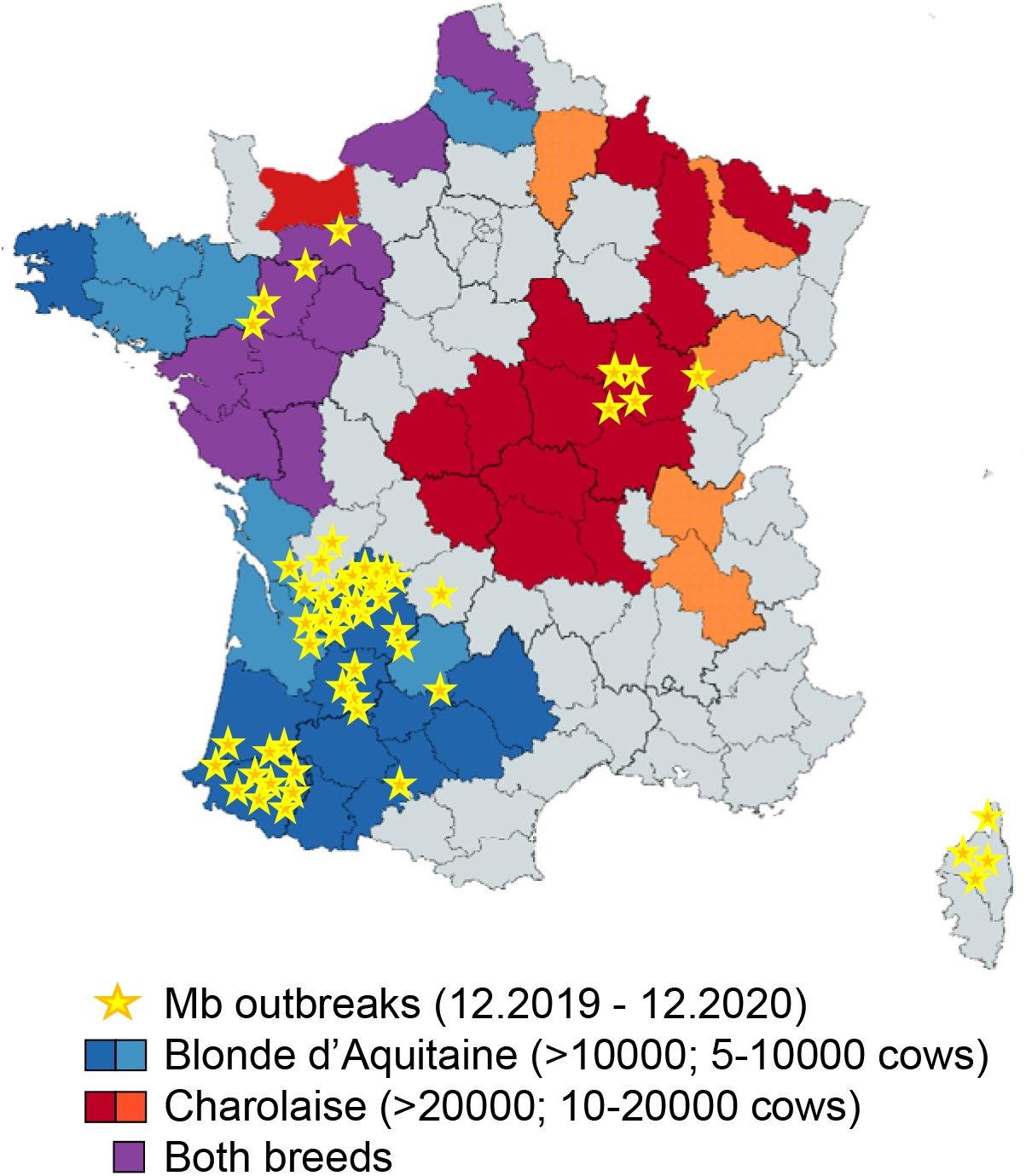
Superposition of Blonde d’Aquitaine and Charolaise beef breeds in French counties where Mb outbreaks were declared between December 2019 and 2020. This map of France shows counties where Mb outbreaks were declared between December 2019 and December 2020 (yellow stars) and was obtained with data extracted from https://www.plateforme-esa.fr/. Herd densities of Blonde d’Aquitaine (blue), Charolaise (red) or both breeds (violet) were extracted from data obtained from https://www.racesdefrance.fr (cows above 3-years-old have been considered).

In addition to the cattle breed, our study pointed towards differences in the host response to distinct mycobacterial strains. Mb strains were better inducers of a lung immune response than Mtb in cattle, in agreement with previous work showing that Mtb H37Rv was attenuated *in vivo* in cattle as compared to Mb AF2122 [13]. *In vitro* studies with bovine AMPs infected with AF2122 or H37Rv revealed differences in the innate cytokine profiles: CCL4, IL-1β, IL-6 and TNFα levels were more elevated in response to AF2122 than H37Rv [8], in agreement with our data. Interestingly, Mb3601, a representative strain of a highly successful genetic cluster that circulates both in cattle and wildlife in France [16] induced an inflammatory signature in the lung more efficiently than Mb AF2122. Whether this correlates with differences in Mb virulence in cattle or other mammals remains to be shown; but if this were the case, the PCLS model would be a practical tool to study and compare the virulence of Mb field strains, as compared to *in vivo* experimental infection of cattle. Contrary to Mtb which is mostly restricted to humans, Mb is adapted to sustain across a large host range through repeated cycles of infection and transmission [57; 58]. This remarkable trait is due to pathogen molecular genetic changes [59] that allow adapted bacilli to manipulate the host immune response to establish infection and disease, and ultimately transmit infection to new, susceptible hosts [60; 61]. We observed weaker inflammation in the bovine lung after infection with Mtb as compared to Mb and it will be interesting to compare the ability of Mtb and Mb to induce inflammation in human PCLS obtained post surgery. This latter comparative analysis could give clues on the links between lung innate inflammatory responses and host-adaptation during TB.

Our most striking observation was the Mb-restricted induction of the type I IFN pathway in the bovine lung. This is in agreement with previous studies in bovine AMPs where cytosolic DNA-sensing pathways, in particular RIG-I, were activated after 48h of infection by Mb AF2122 but not Mtb H37Rv [31]. In agreement with our data these authors also demonstrated induction of the RIG-I signaling pathway by Mb in AMPs [62]. Therefore, AMPs contribute to type I IFN signalling in the lung. However, we also noticed differences between PCLS and AMPs in induction of the IFN signature by Mb: for example, CXCL10 was detected in PCLS but not in AMPs in our study, which may be due to the time point used [63]. However, it is also possible that other cells involved in crosstalk with AMPs contributed to CXCL10 production in response to Mb infection. Since CXCL10 has been proposed as a diagnostic biomarker of Mb infection in cattle [64], it will be interesting to better understand how this key mediator is regulated. Type I interferon favors Mb survival and its induction may be a good manipulation strategy for maintenance of infection. This manipulation mechanism, deciphered *in vitro* in murine bone marrow monocyte-derived MPs, involves triggering of autophagy by cytosolic Mb DNA in turn inducing IFNβ production. Autophagy antagonizes inflammasome activation to the benefit of Mb survival [65; 66]. In C57BL/6 mice treated with IFNAR1 blocking Ab and infected with Mb, the recruitment of neutrophils was reduced but the pro-inflammatory profile of MPs was increased, leading to reduced bacillary burden [67]. No impact on T-cells was observed in this *in vivo* model, revealing a role of type I IFN signaling during the innate phase of the host response to infection. Therefore, Mb exploits type I IFN signaling in many ways and this pathway seems an important avenue to better understand Mb virulence. The PCLS model will greatly help to better dissect out this pathway in the lung during bTB. This could lead to new biomarkers to help genomic selection programs for cattle that are more resistant to bTB, as well as new immunostimulation strategies counteracting the type I IFN pathway. This new knowledge will ultimately improve bTB control, a goal which is so greatly needed at the global level [68].

## ACKNOWLEDGMENTS

We thank the staff from the Abattoir du Perche Vendômois for valuable access to and assistance for bovine post-mortem sampling. We thank Dr. Bojan Stokjovic for his assistance for some PCLS experiment. We are very grateful to Gillian P. McHugo for the drawing of the type I interferon pathway with Ingenuity Pathway analysis. This work was supported by the Veterinary Biocontained research facility Network (VETBIONET), the ANR EpiLungCell (grant ANR-17-CE20-0018) and FEDER/Region Centre Val de Loire ANIMALT grant (FEDER convention number EX007516, Region Centre convention number 2019-00134936, research program number AE-2019-1850). Mobilities between France and Irland were supported by the “ONE-TB” project (PHC Ulysses, funded by Campus France and the Irish Research Council), and the Fédération de Recherche en Infectiologie du Centre Val de Loire (FéRI).

## AUTHOR CONTRIBUTIONS

AR designed and did most of the experiments, obtained funding, analyzed data, prepared all figures and wrote the manuscript. NW obtained funding, supervised all aspects of the work, critically analyzed the data and wrote the manuscript. FC performed experiments and prepared the inocula for experimental infections under BSL3 conditions. AC cultured AMPs, performed ELISA and q-RT-PCR. EDD helped for PCLS experiments and revised the manuscript. MLB provided Mb3601 strain and revised the manuscript. AA provided strain Mtb BTB1558. JAB improved the RNA extraction protocol. DD and QM performed multiplex experiments. FA provided Ab and critically reviewed imaging data. PG helped with transcriptomic analysis and revised the manuscript. SVG obtained funding, designed experiments and revised the manuscript. All authors read and approved the manuscript before publication.

## Supplementary table and figures legends

**Supplementary Table S1. Sequences of primers used in this study.** Primers were designed using Geneious software, in intron-spanning regions when possible. The annealing temperature was set at 60°C. Housekeeping genes used as the reference to calculate ΔCT are indicated in the grey boxes.

**Figure S1: Age and geographical origin of cows used in the study** The Charolaise and Blonde d’Aquitaine cows used were between 3- to 11-years-old, and came from 8 different French departments. Two Blonde d’Aquitaine cows came from the same farm in Indre et Loire; et three Charolaise cows from the same farm in Sarthe. All other animals are from distinct farms. Data represent the age of individual animal and the median and interquartile range.

**Figure S2: Transcriptomic signature after infection with different doses of mycobacteria** Bovine PCLS were obtained as described in Fig. 1 and infected with 10^5^, 5 x 10^5^, 10^6^ or 5 x 10^6^ cfu. RNA were extracted 2 dpi after infection and *SDC4*, *CXCL1*, *HIF1* and *OAS1* gene expression were assessed with the Fluidigm Biomark. Individual data and the mean and standard deviation in each group are presented (n=3 Charolaise). The dotted line represent the level of expression in the uninfected group.

**Figure S3: Cytokines/Chemokines in PCLS supernatants.** Protein levels were measured in PCLS supernatant at 2 dpi post infection with Multiplex. Individual data and the median and interquartile range in each group are presented (n=6). * *p*<0.05 (Wilcoxon non parametric test).

**Figure S4: Structure of bovine PCLS under light microscope.** PCLS were observed under a light microscope (enlargement x40 to x200). PCLS contain numerous alveoli and between one to three bronchioles, with thick and wavy epithelium that can be easily recognized (black asterisk). Thin blood vessels (red dotted lines) were localised next to bronchioles and diffused between alveoli. No blood cells remained inside the endothelium (cows were bled out at the abattoir). Alveolar macrophages can be seen inside alveoli (black arrows).

**Supplementary video 1: Internalisation of Mb3601 in alveolar macrophages after PCLS ex vivo infection.** PCLS were fixed at 2 dpi with 10^5^ cfu of Mb3601-GFP recombinant strain, and labeled with anti-pancytokeratine and anti-MHCII antibodies, respectively revealed with APC, and Alexa 555 conjugated secondary Ab. PCLS were transferred on coverslides and mounted with Fluoromount-G™ Mounting Medium, containing DAPI. Z-stack imaging was performed at x63 enlargment with a confocal microscope. 3D images were analyzed with Leica LAS software.

