## Supplemental Table 1 for "Mycobacterial infection of precision cut lung slices reveals that the type 1 interferon pathway is locally induced by Mycobacterium bovis but not M. tuberculosis in different cattle breeds"

| Gene ID | Reference Sequences | Forward | Reverse |
|---|---|---|---|
| ACTB | NM_173979.3 | ACGGGCAGGTCATCACCATC | AGCACCGTGTTGGCGTAGAG |
| C5aR1 | NM_001007810 | ATACCGTCCTTTGTGTTCCG | ATTGTAAGCGTGACCAGCG |
| CASP1 | XM_024975697 | CTCCCACCTGGCAGGAATAC | AGGAGCTGGAAAAGGAGGGA |
| CASP13 | NM_176638.5 | TCCGGACATTCAACAACCGT | ACCCACAATTCCCCACGATT |
| CASP8 | NM_001045970.2 | AATATTGGGGAGCAGCTGGG | AGGCATCCTTGATGGGTTCC |
| CCL2 | NM_174006 | GCTCGCTCAGCCAGATGCAA | GGACACTTGCTGCTGGTGACTC |
| CCL20 | NM_174263 | TTCGACTGCTGTCTCCGATA | GCACAACTTGTTTCACCCACT |
| CCL5 | NM_175827 | CTGCCTTCGCTGTCCTCCTGATG | TTCTCTGGGTTGGCGCACACCTG |
| CCR1 | NM_001077839.1 | ATGTCTTTGTCGGCGAGAGG | TCTGTGGACAGGAAGGGGAA |
| CD14 | NM_174008 | TCCACAGTCCAGCCGACAAC | AACGGCGCTAGACCAGTCAG |
| CD209 | NM_001145756.1 | CACCCTCGACCACTACACAG | TGAAGAAGCCCAGTGAGACG |
| CD83 | NM_001046590.1 | GAAGGGCAGAGAAACCTGAC | AGAGGTGACTGGGAGGAAAG |
| CLEC4A | NM_001191510.1 | GAAGTTACCACCGTGCTTGC | CCTCTGAAGTCATGCTGCGA |
| CLEC6A | NM_001034479.1 | TACCTGGAAGCCGTTTGGTT | CAAGTGAGCTCCCATCCCAA |
| CLEC7A | NM_001031852.1 | AGGCAAGTGTCTTCCCAGC | ACAACAAGGTGGAGCCATCC |
| CX3CR1 | NM_001102558 | TGGCCTTGGGAACTGTCTTC | TGGCTGTTGATGAGGGCAAA |
| CXCL1 | NM_175700.2 | CCAAACCGAAGTCATAGCC | TCAGTTGGCACTAGCCTTGTTTAGC |
| CXCL10 | NM_001046551 | TTCAGGCAGTCTGAGCCTAC | ACGTGGGCAGGATTGACTTG |
| CXCL2 | NM_174299.3 | GTGTCTCAACCCCGCCGCTC | TCCAGATGGCCTTAGGAGGTGG |
| CXCL3 | NM_001046513.2 | AGCGATGCTGCTCCTGCTCCT | CCATGGGAGCTTCAGGGTTGAG |
| CXCL5 | NM_174300.2 | TGTTTAACCACCACACCGGG | AGGTGGCTATCACTTCCACC |
| CXCL8 | NM_173925.2 | TGAAGCTGCAGTTCTGTCAAG | TTCTGCACCCACTTTTCCTTGG |
| CXCR1 | NM_174360 | ACATGGTTGGTGACTCAGTCTT | CGTGCCGCTGTAATTTCCAA |
| CXCR2 | NM_001101285 | ACAGGTGACAAGCCCAGAATC | CGACCAATCCGGCTGTATAA |
| CXCR3 | NM_001011673 | CCACAGGACTTCAGCCTCAA | CGACTGCCACGATGCCATTA |
| DEFB5 | NM_001130761 | TCGTGCTCCTCTTCCTAGTC | GGCACGAGATCGGAATACAG |
| GAPDH | NM_001034034.2 | GGCATCGTGGAGGGACTTATG | GCCAGTGAGCTTCCCGTTGAG |
| HIF1 | NM_174339 | ACCCTGCACTCAACCAAGAA | TGGGACTGTTAGGCTCAGGT |
| IFITM3 | NM_001078141 | CCTGAACATCTGCTCCCTGG | CTCGGAGACTGCTTGAACGA |
| IFNAR1 | NM_174552.2 | TCCTTTGCCACGTGTCAAGT | AGTAGCGTGAGGGAGACAGA |
| IFNB | XM_005209900 | GCTACAGCTTGCTTCGATTC | TGTGCTGGAGCATCTCATAC |
| IFN-γ | NM_174086 | ACCAGGTCATTCAAAGGAGCAT | TCTGCAGATCATCCACCGGA |
| IL10 | NM_174088.1 | GTGATGCCACAGGCTGAGAA | TGCTCTTGTTTTCGCAGGGCAG |
| IL12p35 | NM_174355 | ACAGAAGGCCAGACAAACTC | AGCCAGACAATGCCCATTAG |
| IL12p40 | NM_174356 | CACCAGCAGCTTCTTCATCA | CTTGTGGCATGTGACTTTGG |
| IL13 | NM_174089.1 | CATGGCGCTCTTATTGACCG | AATGAGCTCCTTGAGGGCTG |
| IL15 | NM_174090 | AACAGCGATGCAGTGCTTTC | TCCTCCAGTTCCTCACATTC |
| IL17A | NM_001008412 | GCCCACCTACTGAGGACAAG | GCTGGATGGTGACAGAGTTC |
| IL17C | 617538 | TGACGTCCACCAGCGCTCCATC | CTGGACCAGCGGCACTGAGTTG |
| IL17F | NM_001192082 | CACTCTGGAGGACCACATTG | GAGTTCAGGGTCCTGTCTTC |
| IL17RA | XM_024992765 | GGCTGAACTGCACAGTCAAG | AGCGTCCACTCGATGTGAAC |
| IL17RB | NM_001083467.1 | GTCCCTCCATGGCTGTGAAC | AGCGCCATGTATCTGTCTCC |
| IL17RC | NM_001075178 | TGCCCTGGTTCCTTCTGTCC | AGGCAGAGCACGTCACCATC |
| IL17RE | XM_010817592.3 | CTGGGAGCCACACTGTAGAC | GTCACGGCCATGACCATCTG |
| IL1-alpha | NM_174092 | CTGAAGAAGAGACGGTTGAG | ATGCATTCCTGGTGGATGAC |
| IL1-beta | NM_174093.1 | CTCTCACAGGAAATGAACCGAG | GCTGCAGGGTGGGCGTATCACC |
| IL21 | NM_198832 | GTGGCCCATAAGTCAAGCTC | CGCTCACAGTGTCTCTTTAC |
| IL22 | NM_001098379 | AGGAGCCCTACATCTTCAAC | CTTCGTCACCTGATGGATTC |
| IL23p19 | NM_001205688 | GATGGCTGTGATCCACAAGG | TGGGAATAGGGCTTGGAGTC |
| IL26 | NM_001205424 | CAGAGCAACGATTCCAGAAG | TCTGCCTGAGGCTATGAAAG |
| IL33 | NM_001075297.1 | GATGGTGGCAGTCATCGGAA | GTAGCTCCACAGAGTGCTCC |
| IL4 | NM_173921.2 | GCCACACGTGCTTGAACAAA | CTTGTGCTCGTCTTGGCTTC |
| IL5 | NM_173922.1 | CAAACTGCACAAGGGGATGC | ATCTTTCTCCTCCACACTTCCT |
| IL6 | NM_173923.2 | TGCTGGTCTTCTGGAGTATC | GTGGCTGGAGTGGTTATTAG |
| IRF3 | NM_001029845.3 | GGAAGGATAAGCCCGACCTG | GAGTCCTTGCTGTGGTCCTC |

| Gene ID | Reference Sequences | Forward | Reverse |
|---|---|---|---|
| IRF7 | NM_001105040.1 | AAGTCTACTGGGAGGTGGGG | CCGAAGTCAAAGATGGGCGT |
| ISG15 | NM_174366 | CGCCCAGAAGATCAATGTGC | TCCTCACCAGGATGGAGATG |
| ITGAM | NM_001039957.1 | TTGAGGCGACGATGGAGTTC | ACTTTCACCTGCCCAGCAAT |
| LAP | NM_203435 | TGCTCCTTGCGCTCCTCTTC | CTCCGAGACAGGTGCCAATC |
| LGP2 | NM_001015545.1 | CCCTTCACTGTGCCTGACTT | AGGTTGTACTGGGCATTGCA |
| MD2 | NM_001046517.1 | AATCGTTGGGTCTGCAACTC | GCGCAATGGGAAATTCATGG |
| MIF | NM_001033608 | GCAAGCCGGCACAGTACATC | CCGCGTTCATGTCGCAGAAG |
| MMP2 | NM_174745 | CCAAGGGTACAGCCTGTTCC | GGCCGGTGCCAGTATCAATG |
| MMP9 | NM_174744 | CGTTCCGACGACATGCTCTG | CATTGCCGTCCTGGGTGTAG |
| MUC1 | NM_174115 | CTCTCCAGGCCATGATAGTG | AAGTGACCATGGAGCTTGAC |
| MX1 | NM_173940.2 | GGCCACATCCCCTTGATCAT | CGTACTGGTCCTTGTCCTGG |
| NLRP3 | NM_001102219.1 | CTCAGTGGCAATACCCTGGG | AGCACTGTCCCAACCACAAT |
| NOD1 | NM_001256563.1 | TGGTCACTCACATCCGAAAC | AGGCCTGAGATCCACATAAG |
| NOD2 | NM_001002889 | CCCAGGGGCTCAGAACTAACA | CCTTCATCCTGGACGTGGTTC |
| NOS2 | NM_001076799 | CTTGAGCGAGTGGTGGATGG | ATCTGAGGGCTGGCATAGGG |
| OAS1Z | NM_001029846.2 | CCAATGGTTCTTCTGCCCCT | GGCAGGAGGTGGTCTTTGAT |
| PKR | NM_178109.3 | TTTTCGCCTCCTCCTCATGC | AACGAATACAGGCTCGCAGA |
| PPIA | NM_178320.2 | TCCGGGATTTATGTGCCAGGG | GCTTGCCATCCAACCACTCAG |
| PTGS2 | NM_174445.2 | CATGGGTGTGAAAGGGAGGAA | ATTTGTGCCCTGGGGATCAG |
| PTX3 | NM_001076259 | TGCCTGCATTTGGGTCAAAG | CACGTTCCTAGGGAAATCAC |
| RIG-I | XM_002689480 | TGTGGTGGAAGATGTTGCGA | AGGGGACATTTCTGCAGCAT |
| S100A7 | NM_174596 | CAGCTTGAGCAGGCCATTAC | CGTGGCTGTGGTTGTGATAG |
| S100A8 | NM_001113725 | CTCCCTGATTGACGTCTACC | TCCAGGCCCACCTTTATCAC |
| S100A9 | NM_001046328 | TGACACCCTGATCCAGAAAG | GCCACCAGCATAATGAACTC |
| SAA3 | NM_181016 | CCTCAAGGAAGCTGGTCAAG | TACCTGGTCCCTGGTCATAC |
| SDC4 | XM_025001268 | AGCTTCAGACAGGGCCTTTC | GTCATGAGCGGGGAAGTAGG |
| STAT1 | NM_001077900.1 | CAAAGGAAGCCCCAGAGCCTAT | GCCACTCTTCTGTGTTCACTTAC |
| TAP | NM_174776.1 | GTAGGAAATCCTGTAAGCTGTG | GTGTCTTGGCCTTCTTTTAC |
| TGFB1 | XM_024977949 | CCTGAGCCAGAGGCGGACTAC | GCTCGGACGTGTTGAAGAAC |
| TLR1 | NM_001046504.1 | ACCCTACTCTGAACCTCAAG | GACTGCACACTGGATTTCTG |
| TLR2 | NM_174197.2 | ACTGGGTGGAGAACCTCATGGTCC | ATCTTCCGCAGCTTACAGAAGC |
| TLR3 | NM_001008664.1 | TTTGCCTGGCTTCCACATCT | GGCGTCTCAAGTTGGAAAGC |
| TLR4 | NM_174198.6 | GCATGGAGCTGAATCTCTAC | CAGGCTAAACTCTGGATAGG |
| TLR5 | NM_001040501.1 | TTCCTGCAACCTCACCCAAG | CTGAGATTGGGCAGGTTTCG |
| TLR6 | NM_001001159 | CTCCGGGAGATAGTCACTTC | GGCCCTGGATTCTATTATGG |
| TLR7 | NM_001033761.1 | GCATCTCTCCAGCCTCCTTT | CACACGTTGTCTTTTGGCCC |
| TLR8 | NM_001033937.1 | AATGCCAAGTCCCAGAGTGG | CCAGCAGCAACTCCCTTAGG |
| TLR9 | NM_183081.1 | GACCTGTCCCACAACAAGCT | TGAAGGGCTGGCTGTTGTAG |
| TNF alpha | NM_173966.3 | TCTTCTCAAGCCTCAAGTAACAAGC | CCATGAGGGCATTGGCATAC |
| TSLP | XM_024995349 | AGAGAGCTACCGGAACATCA | GGGCTGGTCTTCACAGTAGA |
