## Supplementary Figure S1 to S4 for "Mycobacterial infection of precision cut lung slices reveals that the type 1 interferon pathway is locally induced by Mycobacterium bovis but not M. tuberculosis in different cattle breeds"

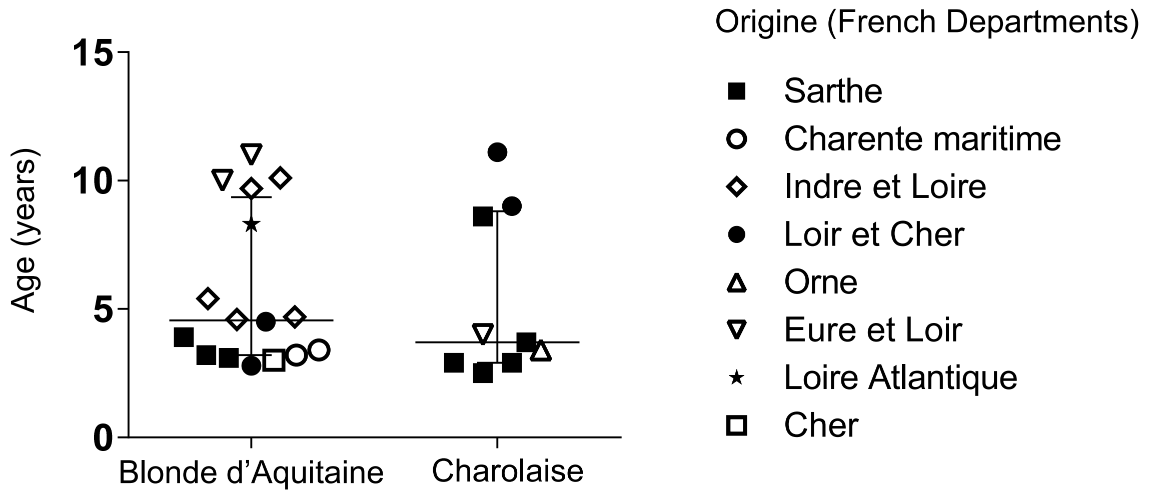


**Figure S1: Age and geographical origin of cows used in the study**

The Charolaise and Blonde d'Aquitaine cows used were between 3- to 11-years-old, and came from 8 different French departments. Two Blonde d'Aquitaine cows came from the same farm in Indre et Loire; et three Charolaise cows from the same farm in Sarthe. All other animals are from distinct farms. Data represent the age of individual animal and the median and interquartile range.

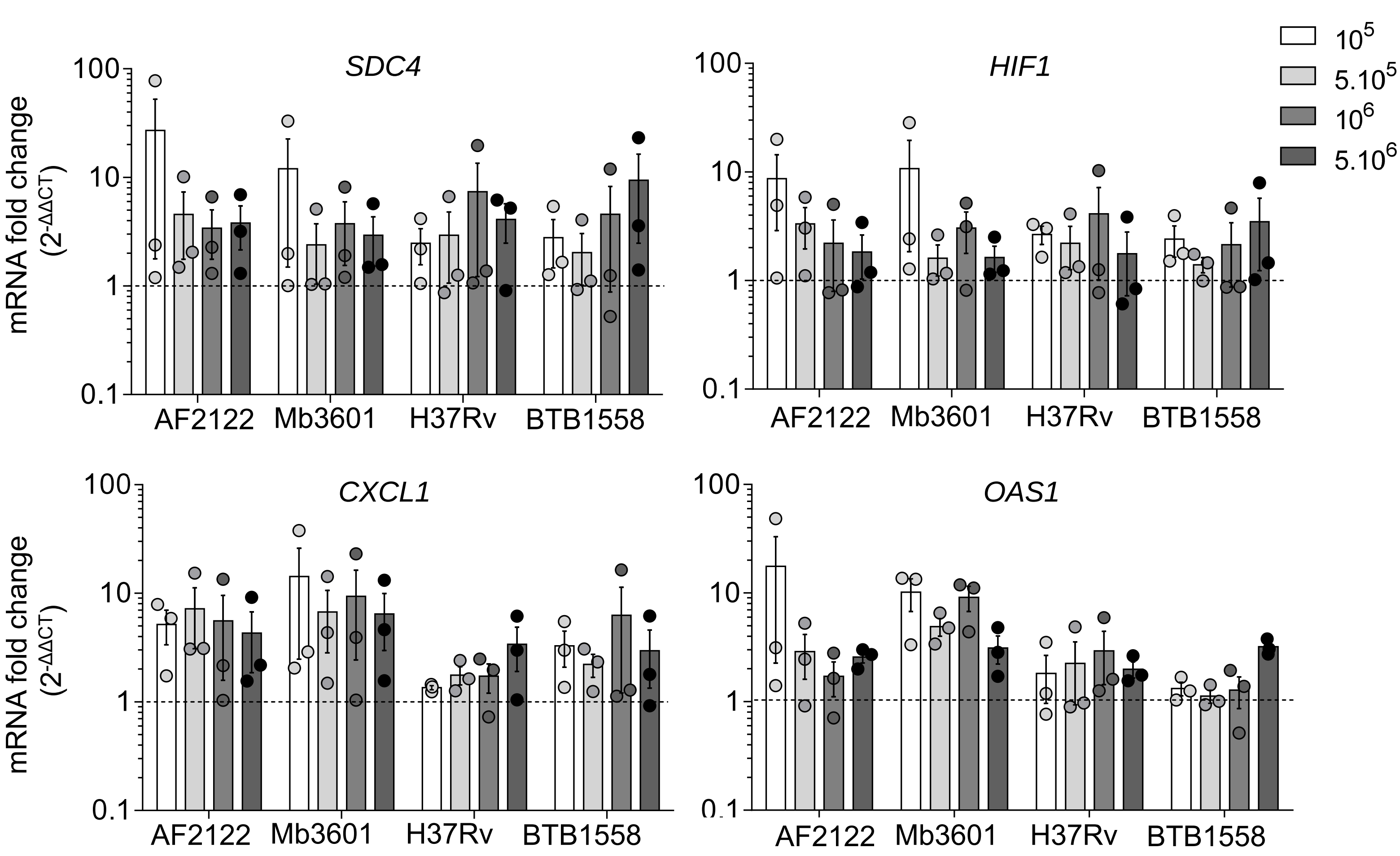


**Figure S2: Transcriptomic signature after infection with different doses of Mycobacteria**

Bovine PCLS were obtained as described in Fig. 1 and infected with $10^5$. 5 x $10^5$. $10^6$ or 5 x $10^6$ cfu. RNA were extracted 2 dpi after infection and *Sdc4*. *Cxcl1*. *Hif1* and *Oas1* gene expression were assessed with the Fluidigm Biomark. Individual data and the mean and standard deviation in each group are presented (n=3 Charolaise). The dotted line represent the level of expression in the uninfected group.

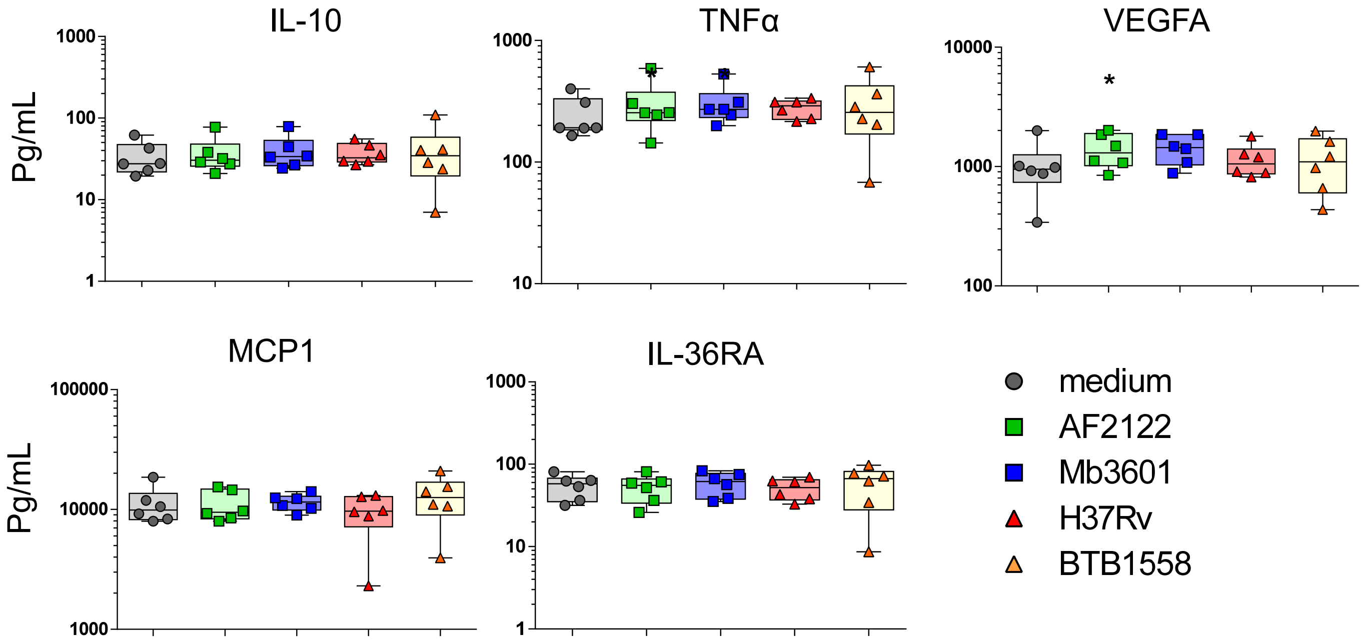


**Figure S3: Cytokines/Chemokines in PCLS supernatants.**
Protein levels were measured in PCLS supernatant at 2 dpi post infection with Multiplex. Individual data and the median and interquartile range in each group are presented (n=6). * $p$<0.05 (Wilcoxon non parametric test).

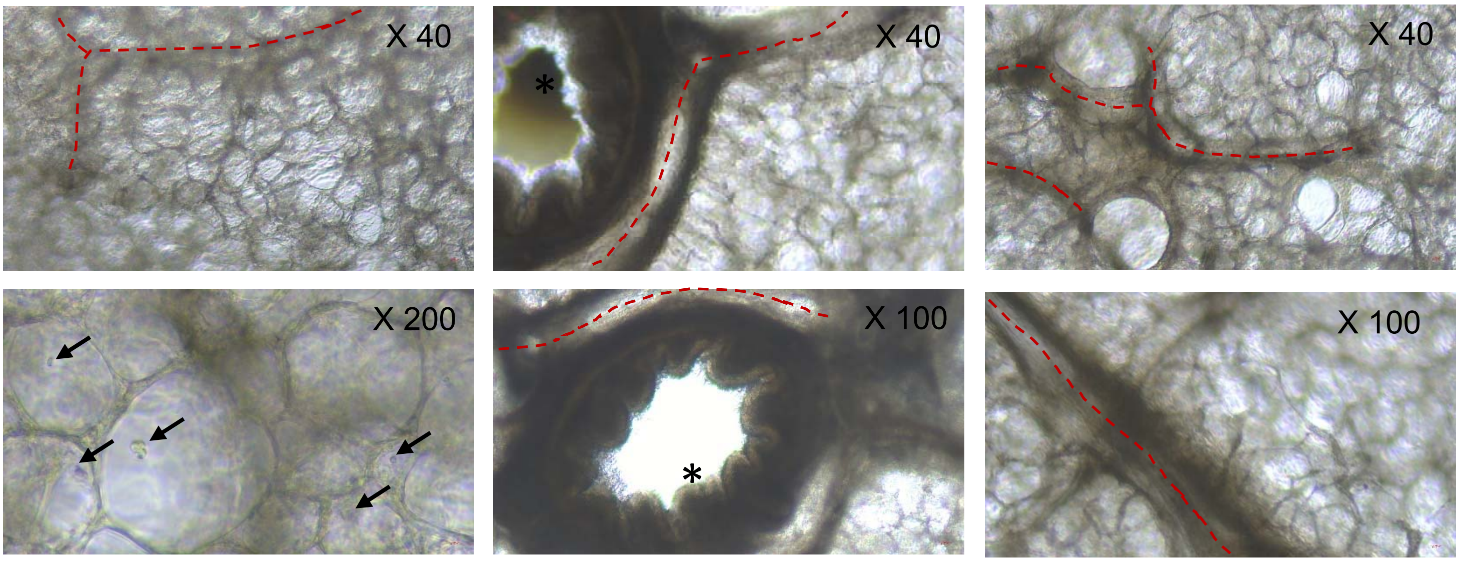


**Figure S4: Structure of bovine PCLS under light microscope.**
PCLS were observed under a light microscope (enlargement x40 to x200). PCLS contain numerous alveoli and between one to three bronchioles, with thick and wavy epithelium that can be easily recognized (black asterisk). Thin blood vessels (red dotted lines) were localised next to bronchioles and diffused between alveoli. No blood cells remained inside the endothelium (cows were bled out at the abattoir). Alveolar macrophages can be seen inside alveoli (black arrows).
